# The F-box protein FBXL-5 governs vitellogenesis and lipid homeostasis in *C. elegans*

**DOI:** 10.1101/2024.04.18.590113

**Authors:** Peter C. Breen, Kendall G. Kanakanui, Martin A. Newman, Robert H. Dowen

## Abstract

The molecular mechanisms that govern the metabolic commitment to reproduction, which often occurs at the expense of somatic reserves, remain poorly understood. We identified the *C. elegans* F-box protein FBXL-5 as a negative regulator of maternal provisioning of vitellogenin lipoproteins, which mediate the transfer of intestinal lipids to the germline. Mutations in *fbxl-5* partially suppress the vitellogenesis defects observed in the heterochronic mutants *lin-4* and *lin-29,* both of which ectopically express *fbxl-5* at the adult developmental stage. FBXL-5 functions in the intestine to negatively regulate expression of the vitellogenin genes; and consistently, intestine-specific over-expression of FBXL-5 is sufficient to inhibit vitellogenesis, restrict lipid accumulation, and shorten lifespan. Our epistasis analyses suggest that *fbxl-5* functions in concert with *cul-6*, a cullin gene, and the Skp1-related gene *skr-3* to regulate vitellogenesis. Additionally, *fbxl-5* acts genetically upstream of *rict-1*, which encodes the core mTORC2 protein Rictor, to govern vitellogenesis. Together, our results reveal an unexpected role for a SCF ubiquitin-ligase complex in controlling intestinal lipid homeostasis by engaging mTORC2 signaling.

## INTRODUCTION

Reproduction is a metabolically expensive process that, in many metazoans, is supported by resources stored in somatic cells. In particular, reallocation of lipids from the soma to the germline, a process called vitellogenesis, is crucial for fertilization, embryonic development, and offspring resiliency (Perez et al., 2017; Perez and Lehner, 2019; Geens et al., 2023; Wang et al., 2023). In the nematode *Caenorhabditis elegans*, synthesis of vitellogenin lipoproteins occurs in the intestine where they mediate the assembly and secretion of triglyceride-rich low density-like lipoprotein (LDL-like) particles that constitute the maternal yolk (Kimble and Sharrock, 1983). The circulating yolk is captured and endocytosed by the RME-2 receptor expressed in mature oocytes (Grant and Hirsh, 1999). Deletion of *rme-2* severely impairs yolk uptake, reduces progeny production by over 20-fold, and extends maternal lifespan (Grant and Hirsh, 1999; Dowen, 2019), which underscores the metabolic trade-off described by the disposable soma theory – that reproductive fitness is inversely related to longevity due to a limited amount of resources that must be balanced between somatic and germline functions (Kirkwood, 1977).

Expression of intestinal vitellogenin lipoproteins is coordinated by a suite of developmental, reproductive, nutritional, and metabolic signals (Perez and Lehner, 2019), which together define the temporal expression and the abundance of vitellogenin lipoproteins. We previously demonstrated that vitellogenesis is regulated by several developmental timing factors that are temporally expressed in the *C. elegans* hypodermis (Dowen et al., 2016). These include the heterochronic genes *lin-4*, *let-7*, and *lin-29*, which when mutated result in reiteration of larval cell divisions and retardation of subsequent developmental events (Chalfie, 1981; Ambros and Horvitz, 1984; Reinhart et al., 2000). Expression of LIN-29, a zinc finger transcription factor, in the hypodermis at the larva-to-adult transition is required for maximal vitellogenin production (Ambros, 1989; Dowen et al., 2016), and mutations that impair *lin-29* expression (*i.e.*, *lin-4* and *let-7*) confer similar vitellogenesis defects. How these factors engage developmental or metabolic signaling pathways in the intestine is poorly understood.

Developmental and environmental regulation of vitellogenesis is coordinated by several pro-growth and homeostatic signaling pathways that function in the intestine. Some of the pathways that control vitellogenin transcription, including the TGF-β and the insulin/insulin-like growth factor signaling (IIS) pathways, are likely to respond to neuronally-secreted ligands (*i.e.*, insulin-like peptides or the TGF-ý family member DBL-1) that are modulated by environmental or nutritional conditions (Goszczynski et al., 2016; Perez and Lehner, 2019). In contrast, other intestinal signaling factors respond to developmental or reproductive signals from hypodermal or germline tissues, respectively (Balklava et al., 2016; Dowen et al., 2016; Goszczynski et al., 2016). We have previously demonstrated that the hypodermal heterochronic pathway engages intestinal mTOR (mechanistic target of rapamycin) signaling to promote vitellogenin gene expression (Dowen et al., 2016). The mTOR kinase resides in two multiprotein complexes, mTOR complex 1 (mTORC1) and mTOR complex 2 (mTORC2), which contain the complex-specific scaffolding proteins DAF-15/Raptor and RICT-1/Rictor, respectively. Loss of either RICT-1 or SGK-1 (serum/glucocorticoid-regulated kinase 1), a downstream target of mTORC2, severely disrupts vitellogenin expression (Dowen et al., 2016). While mTORC2 is necessary for maximal vitellogenin expression, it is more difficult to assess whether mTORC1 is required for vitellogenesis, as complete loss of DAF-15 results in developmental arrest. Although it is well established that insulin/IGF signaling stimulates mTORC2 activity in mammals, it is yet not clear whether mTORC2 is regulated by additional morphogens or pro-growth signaling pathways during development (Oh and Jacinto, 2011).

The factors that mediate crosstalk between the hypodermal heterochronic pathway and the intestinal mTORC2 signaling pathway are unknown. Here, we identify a previously uncharacterized F-box protein, FBXL-5, as an intestinal modulator of vitellogenin expression and intestinal lipid homeostasis in *C. elegans*. Mutations in *fbxl-5* partially suppress the vitellogenin expression defects observed in *lin-29* and *lin-4* mutants, suggesting that FBXL-5 negatively regulates vitellogenesis. This function is performed in concert with the Skp1-related protein SKR-3 and Cullin protein CUL-6, which together form a canonical SCF ubiquitin-ligase complex. Moreover, we demonstrate that FBXL-5 acts within the mTORC2 signaling pathway to regulate intestinal homeostasis. Although FBXL-5 and mTORC2 function together to regulate a broad set of genes, we also found that FBXL-5 acts via a mTORC2-independent mechanism to regulate vitellogenesis and aging pathways. Thus, our work identifies a new cell-autonomous regulator of mTORC2 signaling that couples developmental timing events in the hypodermis to metabolic homeostasis in the intestine.

## MATERIALS AND METHODS

### C. elegans strains

All *C. elegans* strains were cultured at 20**°**C (unless specified otherwise) on NGM media seeded with *E. coli* OP50 as previously described (Brenner, 1974). To synchronize growth of *C. elegans* animals, embryos were isolated by bleaching, washed in M9 buffer, and incubated overnight at room temperature. The resulting synchronized L1s were dropped on seeded plates and grown until adulthood. The stains used in this study are listed in Supplementary Table S1.

### Generation and imaging of transgenic animals

The high-copy *mgIs70[*P*vit-3::GFP]* and single-copy *rhdSi42[*P*vit-3::mCherry]* transgenes have been previously described (Dowen et al., 2016; Torzone et al., 2023). The *fbxl-5* rescue constructs were generated by fusing either the *col-10* promoter (chromosome V: 9,166,416–9,165,291; WS284) or the *vha-6* promoter (chromosome II: 11,439,355–11,438,422; WS284) to a *mCherry::his-58::SL2::fbxl-5* cassette (*fbxl-5* gene: 2,330bp ORF and 176bp of 3’UTR) within the pCFJ151 plasmid via Gibson assembly to generate plasmids pRD99 and pRD98, respectively (Frøkjær-Jensen et al., 2008; Gibson et al., 2009). The sequence-verified plasmids were microinjected into *lin-4(e912); fbxl-5(rhd43); mgIs70[Pvit-3::GFP]* animals at 20 ng/μl, along with 2.5 ng/μl pCFJ90(P*myo-2::mCherry*) and 77.5 ng/μl of 2-Log DNA ladder (New England BioLabs), to generate the DLS327 and DLS330 strains. To generate *rhdIs2* and *rhdIs4*, wild-type animals carrying the extrachromosomal array *Ex[*P*vha-6::mCherry::his-58::SL2::fbxl-5 +* P*myo-2::mCherry]* were gamma-irradiated using a ^137^Cs source (∼3000 rad), resulting in random integration of the array into the genome. The resulting two independent lines were backcrossed at least three times. The *fbxl-5* promoter (chromosome V: 7,746,887 – 7,748,867; WS288) was amplified by PCR and fused to a *mCherry::unc-54 3’UTR* fragment by PCR fusion (Hobert, 2002) to yield a P*fbxl-5::mCherry::unc-54 3’UTR* PCR fragment, which was microinjected into wild-type animals at 75 ng/μl, along with 5 ng/μl P*myo-3::GFP* and 20 ng/μl of 2-Log DNA ladder (New England BioLabs), to generate the DLS344 strain. All strains carrying the *mgIs70* or *rhdSi42* transgenes were imaged with a Nikon SMZ-18 Stereo microscope equipped with a DS-Qi2 monochrome camera, while the DLS344 strain was imaged on a Nikon Eclipse E800 microscope.

### Reporter imaging and quantification

Strains carrying either the *mgIs70* or *rhdSi42* reporter were grown asynchronously, L4s were picked to new plates, and animals were imaged 24 hours later. Alternatively, synchronized animals were grown for 72 hours before imaging. Day 1 adults were mounted on a 2% agarose pad with 25mM levamisole and imaged with a Nikon SMZ-18 Stereo microscope equipped with a DS-Qi2 monochrome camera. The worm bodies were outlined in the brightfield channel and the mean intensities (*i.e.*, gray values) were calculated using the GFP (*mgIs70*) or mCherry (*rhdSi42*) channel using the Fiji 2.14.0 software (Schindelin et al., 2012). Body size measurements (pixels/worm) were simultaneously performed, and the data were converted to mm^2^ based on the known imaging settings. The fluorescence and body size data were plotted in Prism 10 and a one-way ANOVA with a Bonferroni correction was performed to calculate P values.

### CRISPR/Cas9 gene editing

Genomic edits were performed by microinjection of Cas9::crRNA:tracrRNA complexes (Integrated DNA Technologies) into the germline as previously described (Ghanta and Mello, 2020). The crRNA guide sequences are listed in Supplementary Table S2. To generate missense, nonsense, or deletion mutations, single-stranded oligodeoxynucleotides were used as donor molecules. The deletion edits employed two sgRNAs positioned at each end of the deletion site and an oligo that bridged the predicted repair junction. To insert the *3xFLAG::AID* cassette into the 5’ end of the *fbxl-5* gene, dsDNA donor molecules with ∼40 bp homology arms were prepared by PCR using Q5 DNA Polymerase (New England BioLabs) and purified using HighPrep PCR Clean-up beads (MagBio) according to the manufacturers’ instructions. The melted and reannealed DNA repair templates were prepared and microinjected into the germline as previously described (Ghanta and Mello, 2020).

### RNAi Experiments

*E. coli* HT115(DE3) strains carrying plasmids for *fbxl-5* or *lin-29* RNAi knockdown (Ahringer RNAi library), or the control strain containing the empty L4440 plasmid, were grown for ∼19 hrs at 37°C in Luria–Bertani media containing ampicillin (100 μg/ml). The bacteria were concentrated by 20-30x via centrifugation, seeded on NGM plates containing 5 mM isopropyl-β-D-thiogalactoside (IPTG) and 100 μg/ml ampicillin, and maintained at room temperature overnight to induce dsRNA expression. Synchronized L1s were dropped on RNAi plates, grown at 20°C until they were day 1 adults (∼72 hrs), and imaged.

### Lifespan and Brood Size Assays

Animals were grown on standard NGM *E. coli* OP50 plates for at least two generations without starvation prior to lifespan and brood size assays. Longitudinal lifespan assays were performed at 25°C in the presence of 50 μM FUDR as previously described (Torzone et al., 2023). Approximately 150 animals were assayed for each genotype and a log-rank test was used to determine significance. Lifespan assays were repeated at least twice with similar results. For brood size assays (20°C), L4 animals (n=11 per genotype) were picked to individual plates and transferred daily throughout the reproductive period. Progeny were counted as L3, L4s, or as adults. The lowest brood size for each genotype was dropped from the analysis due to potential injury and the data were reported as the mean +/-the standard deviation.

### Oil Red O staining

Animals were fixed in 60% isopropanol and stained for 7 hours with 0.3% Oil Red O exactly as previously described (Torzone et al., 2023). Next, animals were mounted on 2% agarose pads and imaged with either a Nikon SMZ-18 Stereo microscope equipped with a DS-Qi2 monochrome camera (intensity quantification) or a Nikon Ti2 widefield microscope equipped with a DS-Fi3 color camera (representative color images). Quantification of Oil Red O staining was performed in Fiji 2.14.0. Animals were manually outlined, the mean gray value was measured in the worm area, and the value was subtracted from 65,536 (maximum gray value for 16-bit images). The data were plotted in Prism 10 and a one-way ANOVA with a Bonferroni correction was performed to calculate P values.

### Quantitative PCR

Synchronized day 1 adult animals were harvested in M9 buffer, washed, and flash frozen. Isolation of total RNA was performed using Trizol Reagent (Thermo Fisher) with chloroform extraction, followed by isopropanol precipitation. Synthesis of cDNA was performed by oligo(dT) priming using the SuperScript IV VILO Master Mix with ezDNase Kit per the manufacturer’s instructions (Thermo Fisher). Quantitative PCR was performed exactly as previously described (Dowen, 2019). The qPCR primer sequences are listed in Supplementary Table S3. Data from at least three independent experiments were plotted in Prism 10 as the mean fold change relative to wild-type with the standard error of the mean (SEM).

### Western Blot Analyses

Synchronized L1 animals expressing the 3xFLAG::AID::FBXL-5 protein (DLS874 or DLS889) were grown at 20**°**C for either 36 hrs (L3s), 48 hrs (L4s), 72 hrs (day 1 adults), or 96 hrs (day 2 adults) before harvesting them in M9 buffer. The animals were washed three times and pellets were frozen in liquid nitrogen. Whole cell lysates and protein concentrations were determined exactly as previously described (Torzone et al., 2023). The protein samples (50 μg) were resolved by SDS-PAGE, transferred to a PVDF membrane, blocked in 5% nonfat dry milk (BioRad), and probed with either anti-FLAG M2 (F1804, Sigma) or anti-Actin (ab3280, Abcam) antibodies. The developmental time course experiment was performed twice with similar results.

### EMS mutagenesis

The GR2123 strain was mutagenized with ethyl methanesulfonate (M0880, Sigma-Aldrich) exactly as previously described (Dowen et al., 2016). Suppressor mutants displaying increased *mgIs70[Pvit-3::GFP]* reporter expression were selected from ten distinct mutagenesis pools and were singled to individual plates. Mutations that suppressed the egg-laying defects of *lin-4(e912)* were presumed to disrupt *lin-14* function and were not selected for subsequent studies.

Suppressor strains were crossed to the GR2122 strain (*mgIs70[*P*vit-3::GFP]*) and F2 recombinants that displayed the characteristic *egl* and *lon* phenotypes of the *lin-4(e912)* mutant, and that also showed elevated *mgIs70* expression, were singled. Starved animals from the individual plates were then pooled and the genomic DNA was prepared with the Qiagen Gentra Puregene Tissue Kit (Doitsidou et al., 2010). The DNA-Seq libraries were prepared using the NEBNext DNA library preparation kit according to the manufacturer’s instructions (E6040, New England Biolabs) and the libraries were sequenced on an Illumina HiSeq 4000 instrument. Identification of candidate suppressor mutations was performed using in-house scripts according to previously described methods (Minevich et al., 2012).

### mRNA Sequencing

Wild-type and mutant animals were reared on 10 cm NGM agarose plates (5000 animals/plate) until they reached the first day of adulthood. RNA was isolated using Trizol Reagent as described above (see quantitative PCR). The mRNA-Seq libraries were prepared with 1 μg of total RNA using the TruSeq RNA Library Prep Kit v2 per the manufacturer’s instructions (Illumina). Three independent biological replicates were prepared for each condition and libraries were sequenced on an Illumina HiSeq 4000 instrument (single-end, 50bp) at the High Throughput Genomic Sequencing Facility at the University of North Carolina at Chapel Hill. The reads were aligned to the *C. elegans* genome (WS260), and the feature read counts were compiled exactly as previously described (Dowen, 2019). RPKM values and differentially expressed genes (1% FDR) were determined using the DESeq2 algorithm (Love et al., 2014). Tissue-specific enrichment for differential expression was calculated as the number of observed vs. expected differentially expressed genes using previously generated gene lists for each major tissue (Kaletsky et al., 2016). The gene ontology analysis was performed using WormCat (Holdorf et al., 2020). All scatter and violin plots displaying differential expression were generated using the DESeq2 RPKM values and the data were plotted in Prism 10. The raw and processed mRNA-Seq data have been deposited in GEO (GSE248602).

## RESULTS

### Mutations in *T05B11.1/fbxl-5* suppress *lin-4* mutant defects in vitellogenesis

The heterochronic gene *lin-4* encodes a miRNA that governs developmental timing by controlling larval cell divisions in the *C. elegans* hypodermis (Lee et al., 1993). Furthermore, *lin-4* acts non-cell-anonymously to license expression of the vitellogenin genes (*vit-1* through *vit-6*) in the intestine at the L4 larval to adult transition (Dowen et al., 2016). To identify genes that act in the *lin-4* vitellogenesis pathway, we performed an EMS mutagenesis screen to identify mutations that suppress the vitellogenesis defects observed in the *lin-4(e912)* mutant using a P*vit-3::GFP* vitellogenesis reporter as a readout (Supplementary Figure S1A). We selected *lin-4* suppressor mutants that displayed elevated P*vit-3::GFP* expression but that also maintained the egg-laying (*egl*) and long body length (*lon*) defects of the *lin-4(e912)* mutant, which selects against known *lin-4* suppressors (*e.g.*, *lin-14* alleles). After backcrossing to remove unlinked background mutations, we performed whole genome sequencing of the *lin-4* suppressor strains and identified candidate causative mutations using established bioinformatic approaches (Minevich et al., 2012). This resulted in the identification of two putative loss-of-function mutations in the uncharacterized *T05B11.1* gene, which were recovered in two genetically distinct suppressor strains (Supplementary Figure S1B).

The T05B11.1 protein contains a F-box domain at the amino terminus and leucine-rich repeats (LRRs) that span most of the protein (Paysan-Lafosse et al., 2023). Thus, we will subsequently refer to the *T05B11.1* gene as *fbxl-5* (<u>F</u>-<u>b</u>o<u>x</u> leucine rich repeat containing protein 5). F-box proteins canonically function in protein ubiquitination, typically in concert with the Skp1 and Cullin proteins (the SCF complex), which together catalyze E3 ubiquitin ligase activity (Skaar et al., 2013). The F-box domain, and in some cases the LRRs as well, specifically recognizes target substrates and mediates poly-ubiquitination of proteins, resulting in 26S proteasomal degradation (K11/K48 linkages) or altered protein function/localization (K63 linkages). Although there is no clear mammalian orthologue of FBXL-5, highly similar proteins are found in other *Caenorhabditis* species, and it is possible that functional orthologues may exist in other species.

To characterize the role of FBXL-5 in vitellogenesis, we first compared expression levels of the P*vit-3::GFP* reporter between the *lin-4(e912)* single mutant and the *lin-4(e912); fbxl-5(rhd43)* double mutant, finding that while reporter expression is increased in *lin-4(e912)* animals upon loss of *fbxl-5* it does not reach that of wild-type (Figure 1A). Notably, the *rhd43* lesion is a nonsense mutation and is thus predicted to be a strong loss-of-function or null allele. Consistently, knock-down of *fbxl-5* by RNAi only suppressed the P*vit-3::GFP* expression defects observed in the *lin-4* mutant (Supplementary Figure S1C). To rule out the possibility that the high-copy P*vit-3::GFP* reporter lacks the sensitivity to robustly detect the *fbxl-5* suppression phenotype, we employed a single-copy P*vit-3::mCherry* reporter, which closely reflects endogenous *vit-3* levels (Torzone et al., 2023), and generated large deletions within the *fbxl-5* locus using CRISPR/Cas9 genomic editing (Supplementary Table S2). While each of the three *fbxl-5* deletion alleles tested suppressed the *lin-4(e912)* mutant defects in P*vit-3::mCherry* expression, they did not restore levels to those seen in wild-type animals (Figures 1B-C). Interestingly, all *fbxl-5* deletion mutations also reduced the body size of the *lin-4(e912)* mutant (Figure 1D), suggesting that the increase in maternal provisioning may restrict body size or that the FBXL-5 protein regulates additional developmental or metabolic processes outside of vitellogenesis.

**Figure 1.**
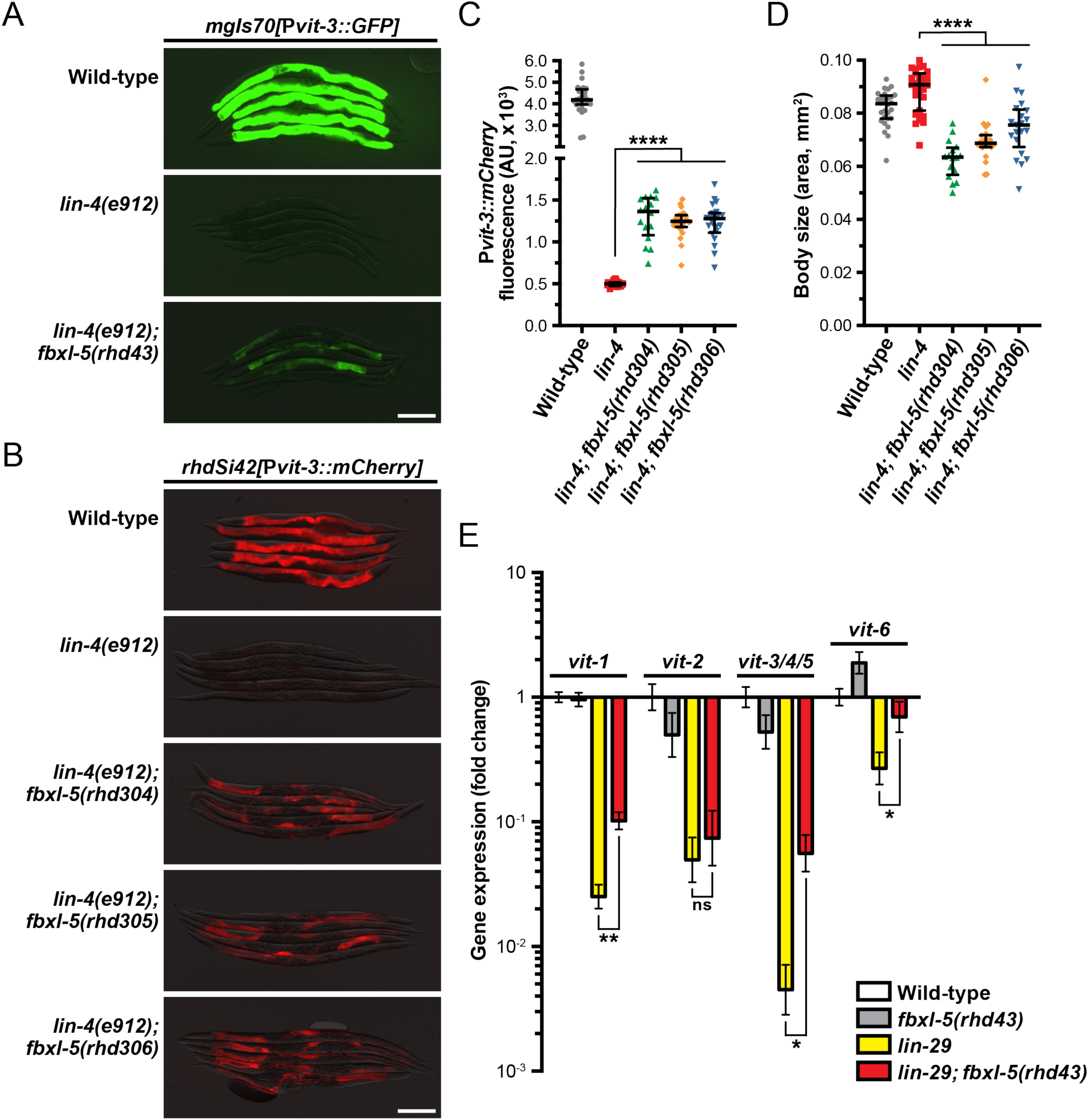
The F-box protein FBXL-5 negatively regulates vitellogenesis in the *C. elegans* heterochronic mutants. Overlaid DIC and fluorescence images of day 1 adult wild-type, *lin-4*, and *lin-4; fbxl-5* animals expressing **(A)** a high-copy P*vit-3::GFP* reporter (*vit-3* promoter fused to *GFP*) or **(B)** a single-copy P*vit-3::mCherry* reporter (scale bars, 200 μm). The *fbxl-5(rhd43)* allele is a nonsense mutation, while the *rhd304-rhd306* are large *fbxl-5* deletion alleles. Quantification of **(C)** P*vit-3::mCherry* fluorescence and **(D)** body size of the strains shown in **(B)** at the day 1 adult stage. Data are displayed as the median and interquartile range (****, P<0.0001, one-way ANOVA). **(E)** Expression of the *vit* genes as measured by RT-qPCR in wild-type, *fbxl-5(rhd43)*, *lin-29(n333)*, and *lin-29(n333); fbxl-5(rhd43)* day 1 adult animals (mean +/-SEM; ns, not significant, *, P<0.05, **, P<0.01, T-test).

Impairment of hypodermal *lin-4* alters the expression of other developmental timing genes resulting in heterochrony. In particular, *lin-4* mutants fail to properly express *lin-29* at the larval to adult transition. Thus, we reasoned that loss of *fbxl-5* may suppress the *vit* expression defects seen in the *lin-29(n333)* mutant. Using RT-qPCR to measure the endogenous *vit* transcripts, we found that mutation of *fbxl-5* also partially suppressed the vitellogenesis defects in *lin-29(n333)* animals; however, loss of *fbxl-5* on its own is not sufficient to increase *vit* transcripts in otherwise wild-type animals (Figure 1E). These data suggest that FBXL-5 may negatively regulate vitellogenesis only when developmental timing is impaired. Together, our results indicate that *fbxl-5* acts genetically downstream or in parallel to *lin-4/lin-29*, possibly in the intestinal cells, to govern vitellogenin expression.

### FBXL-5 governs specific aspects of cellular homeostasis

Although loss of *fbxl-5* only partially suppressed the vitellogenesis defects in the *lin-29* mutant, it’s possible that FBXL-5 plays a broader role in regulating cellular metabolism or lipid allocation in the heterochronic mutants. Taking an unbiased approach, we employed mRNA sequencing (mRNA-Seq) to resolve transcriptome-level differences between the *lin-29* single mutant and the *lin-29; fbxl-5* double mutant. Remarkably, loss of *fbxl-5* had no global impact on the expression levels of the 6,265 genes that are differentially expressed in the *lin-29* mutant (R^2^ = 0.927, Figure 2A). Expression of the *vit-3-5* genes, as well as a handful of other down-regulated genes, did increase in the *lin-29; fbxl-5* double mutant; however, their expression did not return to wild-type. Thus, FBXL-5 has very specific and limited effects on gene expression in the *lin-29* heterochronic mutant.

**Figure 2.**
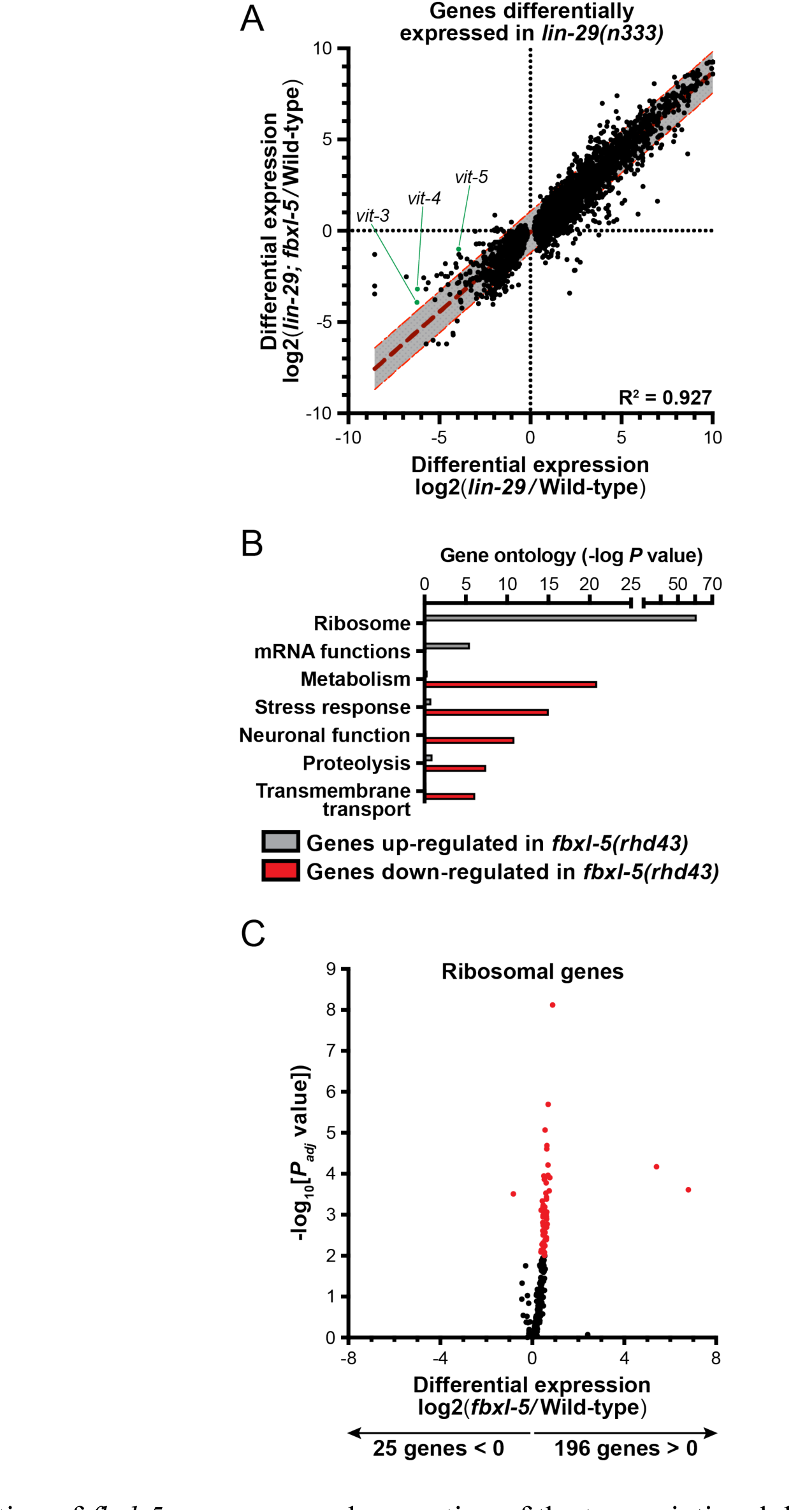
Mutation of *fbxl-5* suppresses only a portion of the transcriptional defects caused by the *lin-29* mutation and prompts a global up-regulation of ribosomal genes. **(A)** A scatter plot of the mRNA-Seq differential expression values comparing the *lin-29(n333)* mutant to the *lin-29(n333); fbxl-5(rhd43)* double mutant for all 6,265 genes differentially expressed in *lin-29(n333)* animals (1% FDR). A linear regression analysis (dark red dashed line) is displayed and the 95% prediction bands for the best-fit line are shown in gray. **(B)** A bar plot showing the P values for the top functional categories derived from a gene ontology analysis of the genes differentially expressed in the *fbxl-5(rhd43)* mutant (1% FDR). **(C)** A volcano plot showing the differential expression values for all annotated ribosomal genes in the *fbxl-5(rhd43)* mutant (red dots indicate differentially expressed genes, 1% FDR).

To further define the role of FBXL-5 in wild-type animals, we performed mRNA-Seq on the *fbxl-5(rhd43)* single mutant. This analysis led to the identification of 828 genes that were differentially expressed upon loss of *fbxl-5*. Mutation of *fbxl-5* resulted in both increased and decreased mRNA levels for genes expressed in each of the major *C. elegans* tissues, which likely represents both the cell-autonomous and non-cell-autonomous effects of mutating *fbxl-5* (Supplementary Figure S2). A gene ontology analysis revealed that the up-regulated genes are enriched in ribosomal and mRNA function, while the down-regulated genes are annotated to act in cellular metabolism, stress response pathways, or proteolysis (Figure 2B). Remarkably, expression of 196 of the 221 *C. elegans* ribosomal genes was increased in the *fbxl-5* mutant (Figure 2C), suggesting that translation is globally upregulated upon loss of *fbxl-5*. Thus, FBXL-5 functions broadly in maintaining cellular energy and protein homeostasis, possibly by governing the activity of a core homeostatic regulator that senses metabolic or developmental signals to tune protein synthesis.

### Heterochronic mutants ectopically express FBXL-5 at adulthood

Since loss of *fbxl-5* in an otherwise wild-type background fails to up-regulate *vit* gene expression (Figure 1E), we reasoned that FBXL-5 levels may be specifically altered in the *lin-4* and *lin-29* heterochronic mutants. To test this hypothesis, we first measured *fbxl-5* transcript levels at the L4 and adult stages in the *lin-4(e912)* mutant by RT-qPCR, finding that *fbxl-5* expression is specifically up-regulated in *lin-4* day 1 adult animals (Figure 3A). Consistently, adult expression of *fbxl-5* is also elevated in the *lin-29* mutant relative to wild-type animals (Supplementary Figure S3A). To measure FBXL-5 protein levels, we inserted a 3xFLAG::AID cassette into the 5’ end of the *fbxl-5* locus by CRISPR/Cas9 editing, which did not disrupt FBXL-5 function and enabled detection by western blotting. We assessed FBXL-5 levels across development by western blot, probing lysates from wild-type and *lin-4(e912)* animals isolated at different developmental stages with an anti-FLAG antibody. Consistent with our gene expression measurements, the *lin-4* mutant ectopically expresses FBXL-5 at the first day of adulthood, which persists at low levels into the second day of adulthood (Figure 3B). We observed a similar FBXL-5 expression pattern for animals subjected to *lin-29* RNAi (Supplementary Figure S3B). Interestingly, wild-type animals have highest expression of FBXL-5 at the L4 stage, suggesting that it may serve a specific role during that developmental stage. Together, these data indicate that heterochronic programs control *fbxl-5* expression, suggesting that the heterochrony of the *lin-4* and *lin-29* mutants may be, at least in part, attributed to improper levels of FBXL-5 during development.

**Figure 3.**
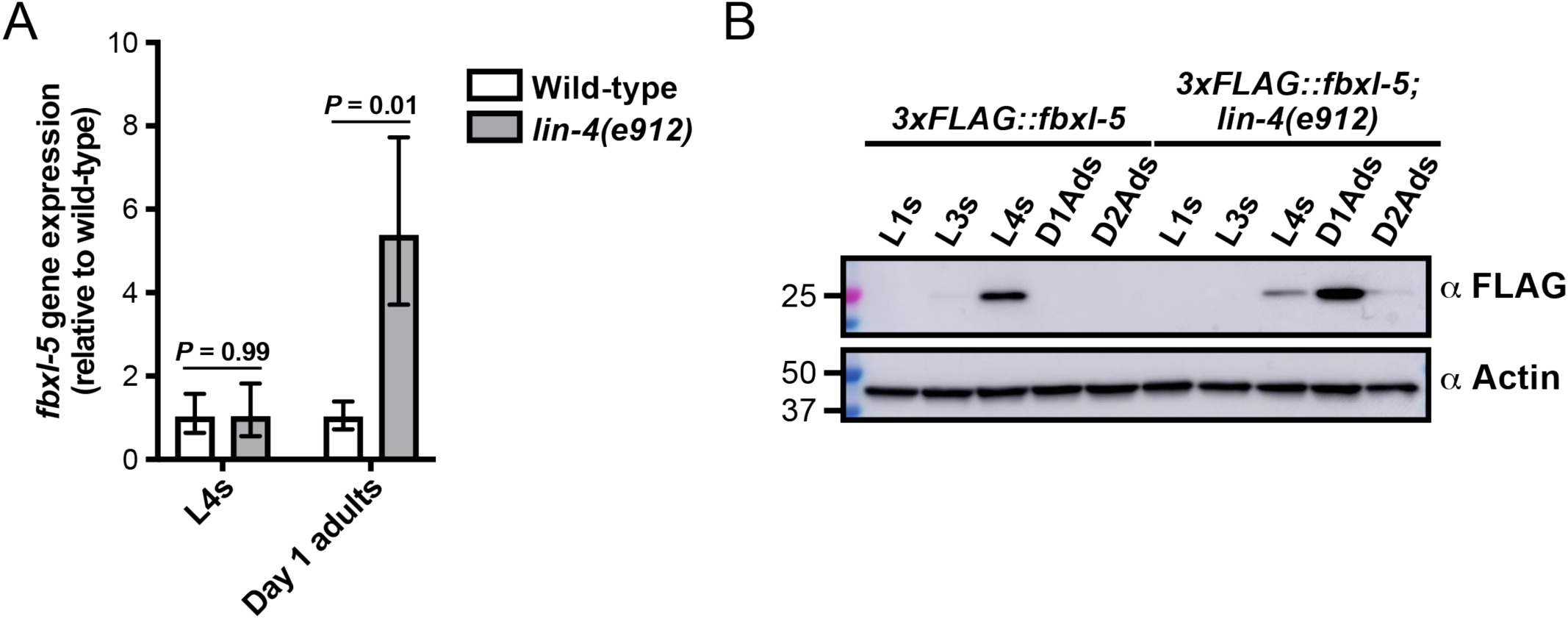
The FBXL-5 protein is misexpressed at the adult stage in the *lin-4* heterochronic mutant. **(A)** mRNA expression levels of *fbxl-5* in wild-type and *lin-4(e912)* animals at the L4 and day 1 adult stage as determined by RT-qPCR (mean +/-SEM; T-test). **(B)** A western blot analysis of lysates from wild-type or *lin-4* mutants expressing an endogenously-tagged 3xFLAG::FBXL-5 protein at the indicated developmental stages. An actin blot is included as a loading control.

### FBXL-5 acts cell-autonomously to govern vitellogenin expression and intestinal homeostasis

Since loss of *fbxl-5* resulted in altered transcription of genes expressed in a variety of tissues, we reasoned that FBXL-5 could act either cell-autonomously or non-cell-autonomously to control vitellogenin expression in the intestine. Importantly, LIN-29 functions in the hypodermis to regulate vitellogenesis in a non-cell-autonomous fashion (Dowen et al., 2016), and it is possible that ectopic expression of *fbxl-5* in the hypodermis is partially responsible for the impaired vitellogenin production that is observed in the *lin-29* mutant. To investigate the tissue-specificity of FBXL-5, we first generated a transgenic strain expressing a P*fbxl-5::mCherry* transcriptional reporter, comprised of a ∼2kb fragment of the *fbxl-5* promoter driving the *mCherry* gene, and inspected animals for mCherry expression by fluorescence microscopy. This analysis revealed that *fbxl-5* is indeed weakly expressed in the intestine of wild-type young adults; however, it is also expressed in small set of neurons in the head and in hypodermal and/or body muscle cells (Figure 4A), which is consistent with the widespread transcriptional changes that we observed. We reasoned that *fbxl-5* likely acts in the intestine or in the hypodermis (the site of *lin-29* action) to regulate vitellogenesis. Thus, we generated intestine-specific and hypodermis-specific *fblx-5* rescue constructs to test whether the *fblx-5(rhd43)* mutation could be rescued by tissue-specific expression of *fblx-5*. Rescue of *fblx-5* in the intestine, but not in the hypodermis, reversed the suppressive effects of *fblx-5(rhd43)* in the *lin-4* mutant background back to baseline P*vit-3::GFP* levels (Figure 4B-C). Surprisingly, hypodermal *fblx-5* over-expression further enhanced the suppressive effects of the *rhd43* allele, suggesting that FBXL-5 might promote vitellogenin expression when expressed outside of the intestine. While the vitellogenesis phenotype was rescued by intestinal expression of *fbxl-5*, the small body size phenotype that we observed in the *lin-4(e912)*; *fblx-5(rhd43)* double mutant was not rescued by intestinal or hypodermal *fbxl-5* expression (Figure 4D), suggesting that FBXL-5 may act non-cell-autonomously to control body size.

**Figure 4.**
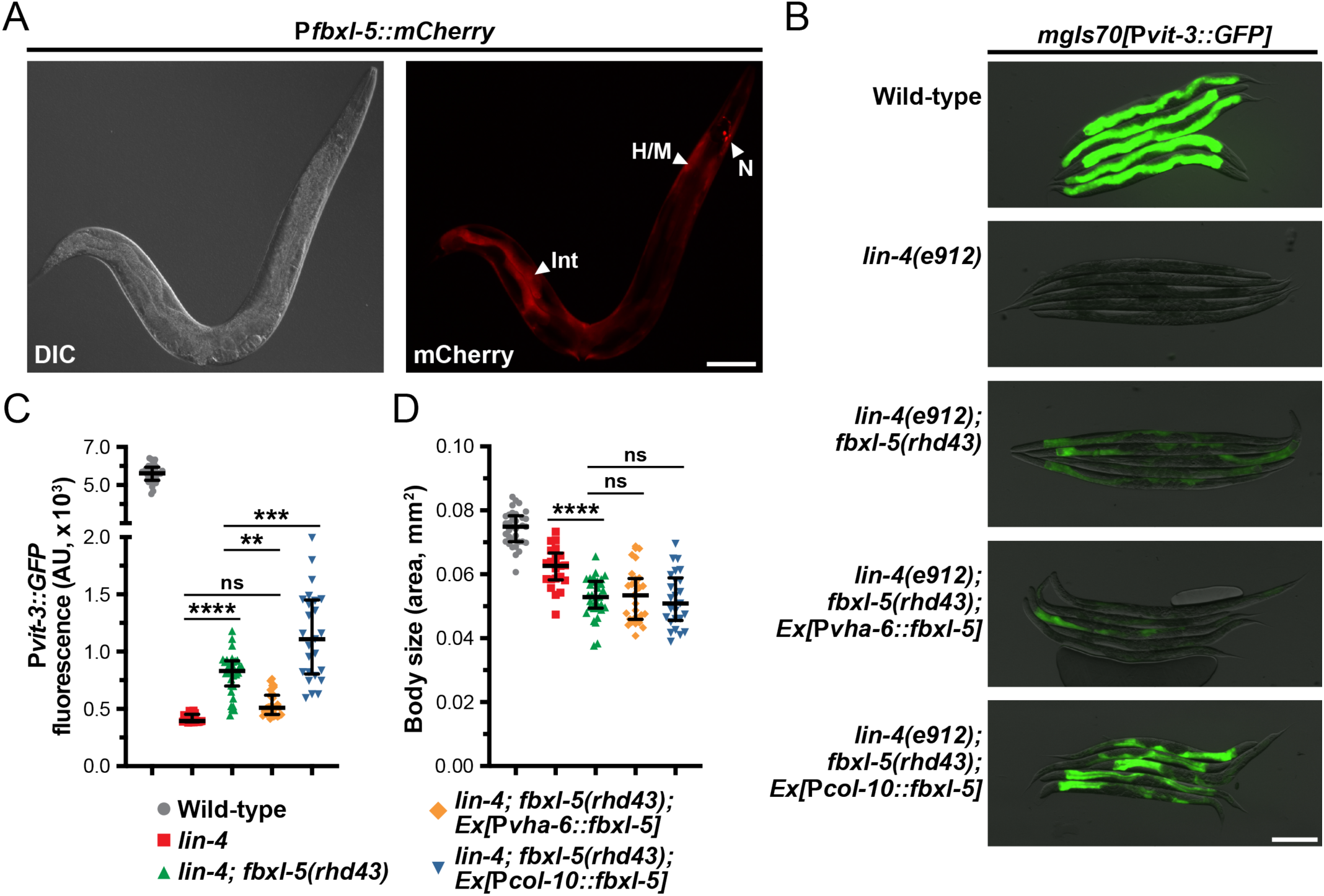
The FBXL-5 protein acts cell-autonomously to control vitellogenesis. **(A)** A representative DIC and mCherry fluorescence image of an animal expressing a P*fbxl-5::mCherry* transcriptional reporter (Int, intestine; H, hypodermis; M, muscle; N, neurons; scale bar, 100 μm). **(B)** Representative overlaid DIC and P*vit-3::GFP* fluorescence images (scale bar, 200 μm), **(C)** P*vit-3::GFP* reporter quantification, and **(D)** body sizes of day 1 adult wild-type, *lin-4(e912)*, and *lin-4(e912); fbxl-5(rhd43)* animals, as well as the indicated *lin-4(e912); fbxl-5(rhd43)* tissue-specific rescue strains (P*vha-6*, intestinal-specific rescue; P*col-10*, hypodermal-specific rescue). **(C, D)** Data are plotted as the median and interquartile range (ns, not significant, **, P<0.01, ***, P<0.001, ****, P<0.0001, one-way ANOVA).

Since *fbxl-5* is ectopically expressed in the *lin-4* and *lin-29* heterochronic mutants, we reasoned that the consequences of FBXL-5 misexpression should be mirrored by constitutive over-expression of *fbxl-5* in the intestine of otherwise wild-type animals. Thus, we integrated a P*vha-6::fbxl-5* extrachromosomal array into the genome via gamma irradiation and crossed the resulting strains to our vitellogenesis reporter. Consistent with our *fbxl-5* rescue experiments, constitutive over-expression of *fbxl-5* in the intestine is sufficient to abrogate P*vit-3::GFP* expression (Figure 5A-B). Furthermore, *fbxl-5* over-expression prevented the accumulation of intestinal lipids and restricted body size (Figure 5C-D), suggesting that FBXL-5 might impair nutrient absorption or stimulate lipid catabolism pathways. Lipid levels are reduced both at the L4 larval stage and during reproduction by *fbxl-5* over-expression (Figure 5E), suggesting that FBXL-5 does not act exclusively on a larval-specific metabolic regulator.

**Figure 5.**
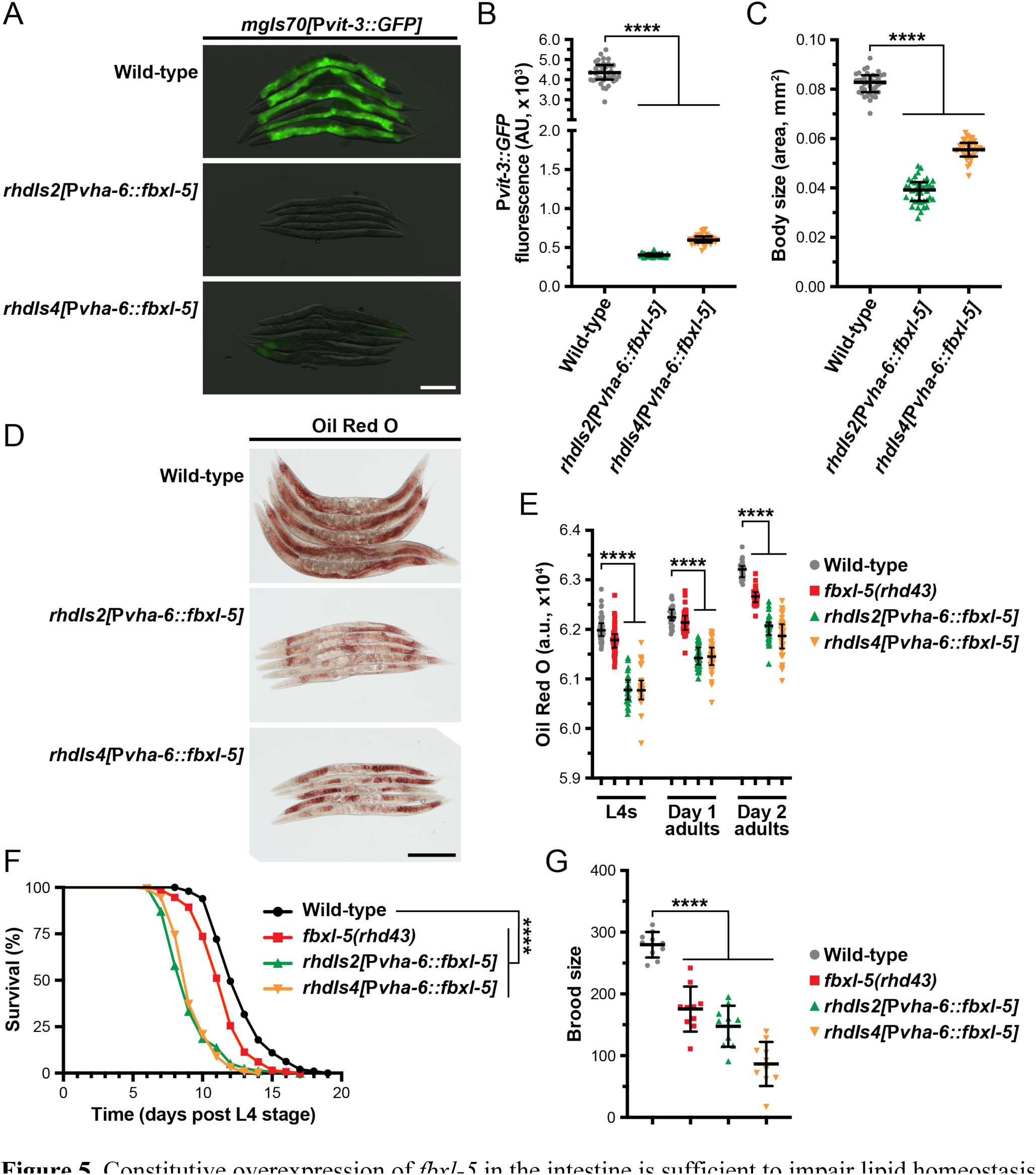
Constitutive overexpression of *fbxl-5* in the intestine is sufficient to impair lipid homeostasis and shorten lifespan. **(A)** Overlaid DIC and P*vit-3::GFP* fluorescence images (scale bar, 200 μm), **(B)** P*vit-3::GFP* reporter quantification, **(C)** body size, and **(D)** representative images (scale bar, 200 μm) of Oil Red O-stained wild-type or *fbxl-5* intestinal overexpression animals (P*vha-6::fbxl-5*, two independently integrated array lines). **(E)** Quantification of Oil Red O staining for wild-type, *fbxl-5(rhd43)*, and *fbxl-5* overexpression animals at the L4, day 1 adult, and day 2 adult stages reveals that FBXL-5 restricts lipid accumulation. **(F)** Longitudinal lifespans (25**°**C with FUDR; ****, P<0.0001, Log-rank test) and **(G)** brood size (20**°**C; mean +/-SD; ****, P<0.0001, one-way ANOVA) of wild-type, *fbxl-5(rhd43)*, and *fbxl-5* overexpression animals. **(B, C, E)** Data are plotted as the median and interquartile range (****, P<0.0001, one-way ANOVA).

Impaired vitellogenin production can indirectly extend organismal lifespan by stimulating intestinal autophagy and lipid catabolism (Murphy et al., 2003; Seah et al., 2016). Thus, we tested whether *fbxl-5* over-expression, which severely reduces vitellogenin synthesis, increases longevity. Surprisingly, over-expression of *fbxl-5* in the intestine reduced the lifespan of otherwise wild-type animals (Figure 5F), which is consistent with FBXL-5 playing a more widespread role in metabolic regulation. Moreover, the *fbxl-5(rhd43)* mutant also displayed a modest, but statistically significant, reduction in lifespan, which is consistent with the modest reduction in lipid levels and brood size that we observed in these mutant animals (Figure 5E and 5G). Predictably, intestinal over-expression of *fbxl-5* reduced progeny production (Figure 5G), likely a consequence of the impaired vitellogenin production. Together, these data suggest that FBXL-5 can function broadly in the intestine to alter metabolic, reproduction, and longevity programs.

### FBXL-5 functions in a canonical SCF complex

One canonical function of F-box proteins is to mediate poly-ubiquitination and proteasomal degradation of their substrates, which requires coordination with the Skp1 and Cullin proteins. While *C. elegans* expresses ∼520 F-box proteins, the genome only contains 21 Skp1-related (*skr*) genes and 6 Cullin (*cul*) genes, suggesting that these factors have less functional redundancy than the F-box family (Nayak et al., 2002; Thomas, 2006). To determine if FBXL-5 functions in concert with the SKR and CUL proteins, we performed a small-scale RNAi screen to identify the *skr*/*cul* genes that are required for reducing P*vit-3::GFP* reporter expression in *fbxl-5* over-expression animals. Knock-down of *skr-3* by RNAi modestly increased GFP levels in the reporter strain, which was recapitulated using the *skr-3(ok365)* mutant allele (Figure 6A). Although we also identified *cul-6* as a candidate in our RNAi screen, the *cul-6(ok1614)* mutation failed to replicate this finding; however, mutation of both *skr-3* and *cul-6* strongly increased P*vit-3::GFP* expression in *fbxl-5* over-expression animals relative to the control (Figure 6A and 6B). Further mutation of *skr-4* or *skr-5*, which both share significant similarity to *skr-3* (Nayak et al., 2002), failed to dramatically enhance the *skr-3(ok365); cul-6(ok365)* double mutant. Together, these data suggest that FBXL-5 functions in concert with SKR-3 and CUL-6 to restrict vitellogenin production; however, it is likely that FBXL-5 acts with other redundantly-acting factors, as P*vit-3::GFP* expression levels are only partially restored to wild-type levels in *skr-3; cul-6* double mutant.

**Figure 6.**
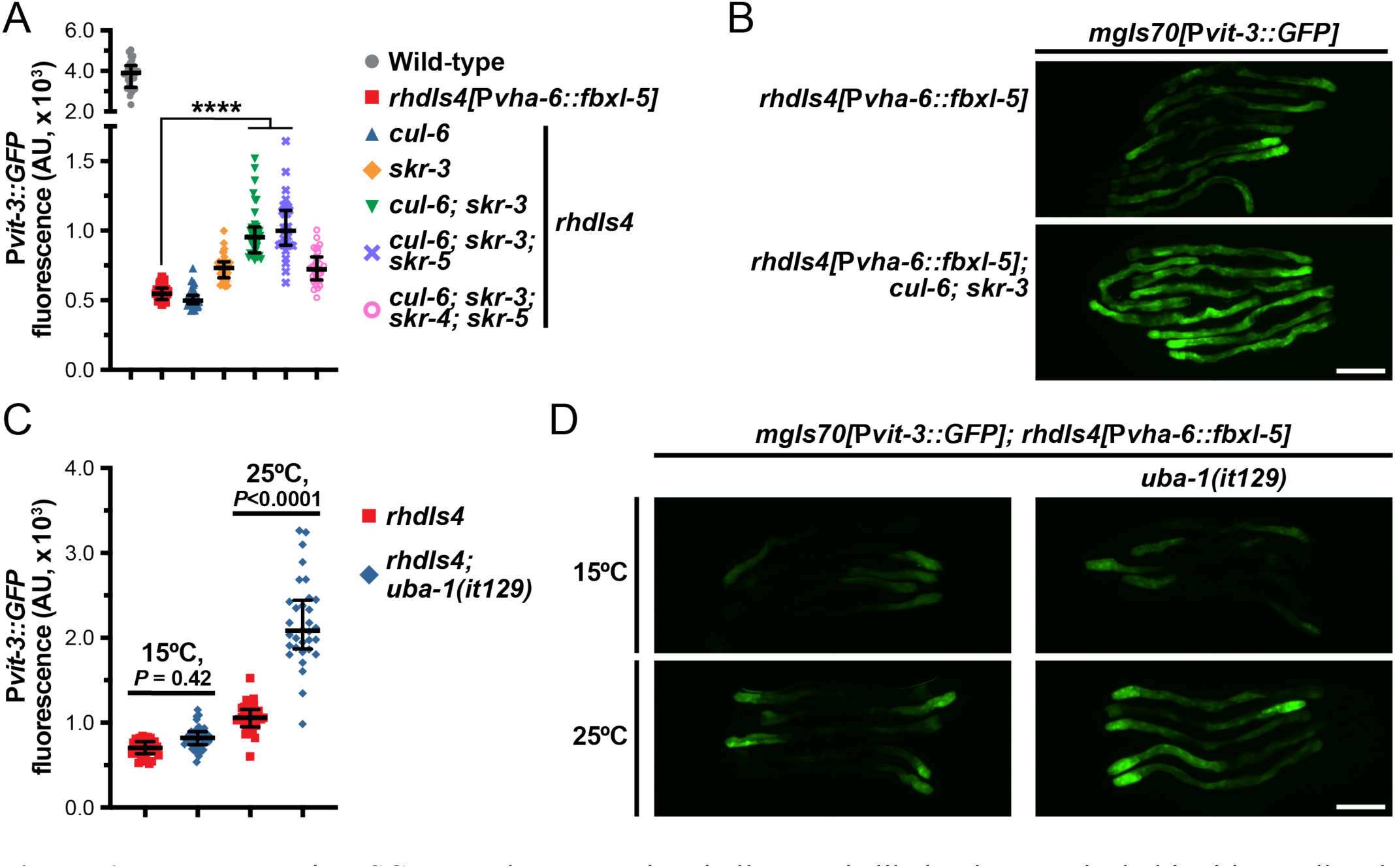
FBXL-5 acts in a SCF complex to restrict vitellogenesis likely via canonical ubiquitin-mediated proteasomal degradation. **(A)** P*vit-3::GFP* fluorescence quantification and **(B)** representative P*vit-3::GFP* images of *fbxl-5* intestinal overexpression animals (*rhdIs4*) carrying the indicated *skr* (Skp1 related) or *cul* (<u>Cul</u>lin) mutations suggesting that SKR-3 and CUL-6 act with FBXL-5 in a SCF complex. **(C)** P*vit-3::GFP* fluorescence quantification and **(D)** representative P*vit-3::GFP* images of *rhdIs4* or *rhdIs4*; *uba-1(it129)* animals. The *uba-1(it129)* mutation is a temperature sensitive loss-of-function allele at 25**°**C. **(A, C)** Data are plotted as the median and interquartile range (****, P<0.0001, one-way ANOVA). **(B, D)** Scale bars are 200 μm.

To further examine the role of poly-ubiquitination in the FBXL-5-dependent regulation of vitellogenesis, we introduced the *uba-1(it129ts)* temperature-sensitive mutation into the *fbxl-5* over-expression strain. UBA-1 is the sole E1 ubiquitin-activating enzyme in *C. elegans* and its activity is required for the ubiquitin proteolytic pathway (Kulkarni and Smith, 2008). Consistent with our previous results, the *uba-1(it129ts)* mutation partially suppressed the effects of *fbxl-5* over-expression at the restrictive, but not permissive, temperature (Figure 6C and 6D). Together, our results suggest that FBXL-5 acts in a SCF complex with SKR-3 and CUL-6 to promote poly-ubiquitination-mediated proteasomal degradation.

### FBXL-5 acts in concert with mTORC2 to govern intestinal homeostasis

We have previously shown that hypodermal *lin-29* acts genetically upstream of intestinal mTORC2 signaling to regulate vitellogenesis (Dowen et al., 2016). We hypothesized that FBXL-5 might be responsible for modulating the signaling between LIN-29 and mTORC2, which we tested by performing genetic epistasis analyses. While knock-down of *fbxl-5* by RNAi increased P*vit-3::GFP* expression 3.3-fold in *lin-4(e912)* and 3.9-fold in *lin-29(n333)* mutant animals, the suppressive effects seen in *rict-1(mg360)* and *sgk-1(ok538)* animals were less striking (1.6- and 0.9-fold, respectively; Figure 7A and Supplementary Figure S4A). The *mg360* allele is a missense mutation in *rict-1*, and thus, this mutant could retain some *rict-1* function, which complicates the interpretation of our epistasis analysis.

**Figure 7.**
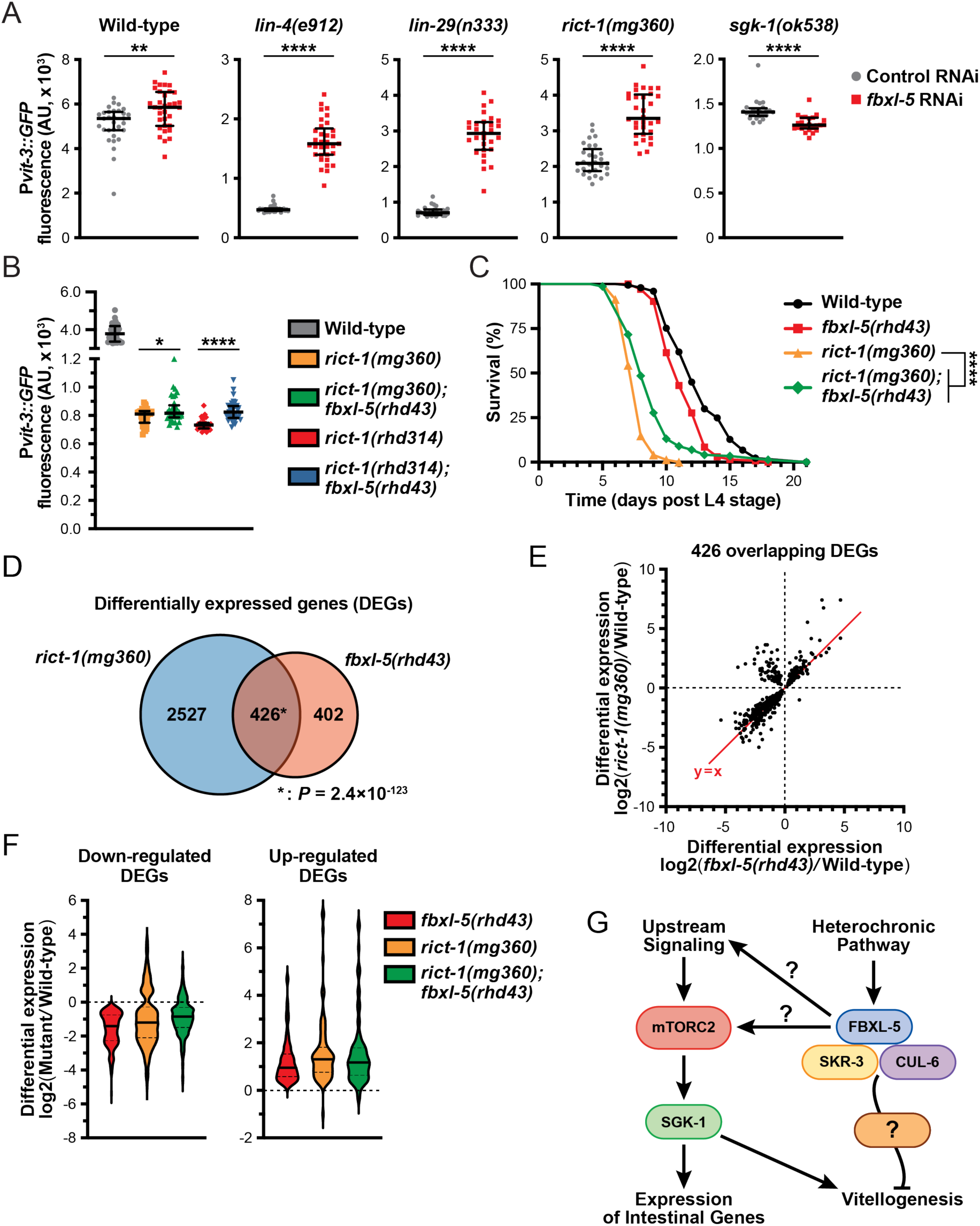
FBXL-5 functions in the mTORC2 pathway. **(A)** Quantification of P*vit-3::GFP* fluorescence in the indicated heterochronic (*lin-4*, *lin-29*) or mTORC2 (*rict-1*, *sgk-1*) mutants following treatment with either control or *fbxl-5* RNAi (median and interquartile range; ****, P<0.0001, T-test). **(B)** P*vit-3::GFP* fluorescence quantification of strains carrying either the *rict-1(mg360)* missense mutation or the *rict-1(rhd314)* null mutation, which lacks the entire *rict-1* locus (median and interquartile range; *, P<0.05, ****, P<0.0001, T-test). The *fbxl-5(rhd43)* mutation modestly suppresses both *rict-1* mutations. **(C)** Longitudinal lifespan assays of the indicated strains reared at 25**°**C in the presence of FUDR demonstrating that the *fbxl-5(rhd43)* mutation extends the lifespan of short-lived *rict-1(mg360)* animals (****, P<0.0001, Log-rank test). **(D)** A Venn diagram showing the overlap of genes differentially expressed in the *rict-1(mg360)* and *fbxl-5(rhd43)* mutants relative to wild-type (1% FDR; *, hypergeometric P value). **(E)** A scatter plot of the differential expression values of the 426 genes co-regulated by RICT-1 and FBXL-5 indicating that most genes are similarly regulated. (F) Violin plots comparing the differential expression values of the 426 RICT-1/FBXL-5 co-regulated genes for the *fbxl-5(rhd43)* and *rict-1(mg360)* single mutants, as well as the *rict-1(mg360)*; *fbxl-5(rhd43)* double mutant, showing that loss of both genes fails to produce an additive effect. **(G)** A proposed model of FBXL-5 action in the *C. elegans* intestine.

Therefore, we generated a complete *rict-1* deletion mutant by CRISPR/Cas9 editing, removing the entire sequence between the start and stop codons, including the *pqn-32* gene that is positioned between exons 10 and 11, and measured P*vit-3::GFP* reporter expression. Consistent with the *mg360* allele, complete deletion of *rict-1* resulted in a robust decrease in P*vit-3::GFP* levels, while further mutation of *fbxl-5* resulted in a modest, but significant, 1.1-fold increase in reporter expression (Figure 7B). This weak suppression phenotype was not limited to vitellogenesis, as the *fbxl-5(rhd43)* mutation also partially suppressed the short lifespan of the *rict-1(mg360)* mutant (Figure 7C). To eliminate the possibility that FBXL-5 non-specifically restricts vitellogenin expression in the intestine, we performed *fbxl-5* RNAi on *daf-2(e1370)* mutants, which also have reduced P*vit-3::GFP* reporter expression (Dowen et al., 2016). Knock-down of *fbxl-5* had no impact on vitellogenin expression in *daf-2* mutant animals (Supplementary Figure S4B). Together, our results suggest that *fbxl-5* acts genetically upstream of *rict-1* and *sgk-1* to regulate vitellogenesis; however, it’s also likely that *fbxl-5* acts simultaneously in a pathway parallel to *rict-1* to regulate lifespan and vitellogenesis via a mTORC2-independent mechanism.

To further define the relationship between FBXL-5 and RICT-1 signaling, we performed mRNA-Seq on the *rict-1(mg360)* mutant and compared the differentially expressed genes (DEGs) between the *rict-1* and *fbxl-5* mutants. There was a striking overlap between these datasets, with 51% of the *fbxl-5* DEGs also misexpressed in the *rict-1* mutant (Figure 7D). The vast majority of these overlapping DEGs showed similar directionalities and degrees of mis-expression between the two mutants; however, a small cluster of genes were up-regulated in the *rict-1(mg360)* mutant but down-regulated in *fbxl-5(rhd43)* animals (Figure 7E). This could be a result of FBXL-5 having different roles within the intestine or having unique functions in different tissues. With many genes similarly altered in *rict-1* and *fbxl-5* mutants, we performed mRNA sequencing on the *rict-1(mg360); fbxl-5(rhd43)* double mutant and calculated the differential expression values for the 426 co-regulated genes. Simultaneous loss of both *rict-1* and *fbxl-5* did not further alter gene expression levels compared to the single mutants (Figure 7F), suggesting that these genes are regulated by FBXL-5 and RICT-1 in a linear pathway. Together, our results indicate that FBXL-5 regulates gene expression via mTORC2-dependent and independent mechanisms (Figure 7G).

## DISCUSSION

Here, we demonstrate that the previously uncharacterized FBXL-5 protein functions in the mTORC2 pathway to govern lipid homeostasis. While much of FBXL-5’s function aligns with the hypothesis that it is a positive regulator of mTORC2 signaling, we have also shown that FBXL-5 acts in a mTORC2-independent pathway to confer modest effects on vitellogenesis and organismal lifespan (Figure 7G). It is likely that FBXL-5 functions in concert with SKR-3 and CUL-6 to mediate poly-ubiquitination-dependent proteasomal degradation of at least one target protein; however, it is not surprising that loss of *fbxl-5* yields pleiotropic effects, as it is also equally possible that FBXL-5 acts on multiple targets and/or pathways. Alternatively, the effects of loss of *fbxl-5* on mTORC2 signaling may have an indirect effect on other pathways, including the mTORC1 pathway. Notably, the mTORC1 complex shares many of the same proteins as mTORC2, which may be redistributed to mTORC1 upon loss of FBXL-5 (Nukazuka et al., 2011). Nonetheless, our results demonstrate a role for a FBXL-5-containing SCF complex in the regulation vitellogenesis, which has not been previously described.

Mutation of the *lin-4* gene results in heterochrony due to cell lineage defects primarily in the larval hypodermis (Chalfie, 1981; Ambros and Horvitz, 1984). While there is significant evidence that *lin-4* functions in a cell-autonomous fashion in several tissues (Zhang and Fire, 2010), loss of *lin-4* or *lin-29* impairs intestinal vitellogenin gene expression in a non-cell-autonomous manner (Dowen et al., 2016). This response is partially governed by intestinal FBXL-5. Consistent with these observations, migration of hermaphrodite-specific neurons is controlled non-cell-autonomously by hypodermal factors, which promote the expression of receptors or cell-adhesion molecules that maintain neuron-hypodermal contacts (Pedersen et al., 2013; Salzberg et al., 2013). High levels of FBXL-5, as modeled in the *fbxl-5* over-expression animals, could impair communication between hypodermal and intestinal cells by restricting an inter-tissue signal pathway that in turn couples to intestinal mTORC2 signaling. The *lin-4* and *lin-29* mutants both display ectopic expression of *fbxl-5* in adults, which could be inhibiting an adult-specific signal from the hypodermis that is required for initiating vitellogenesis; however, additional studies are needed to resolve what types of developmental signals are released from the hypodermis to coordinate intestinal metabolism.

The *C. elegans* genome encodes ∼520 F-box-containing proteins that serve as adaptors for SCF-mediated, ubiquitin-dependent proteolysis. In contrast, the human genome encodes ∼68 F-box proteins (Jin et al., 2004), suggesting that F-box proteins may serve redundant or additional roles in *C. elegans*, which may include responding to foreign or pathogenic proteins that nematodes commonly encounter in the wild (Thomas, 2006), performing nematode-specific developmental or reproductive functions (Nayak et al., 2002; Fielenbach et al., 2007; Gao et al., 2008), or providing tissue-specific responses to a ubiquitous signaling molecule. Importantly, only a handful of F-box proteins have clearly defined roles in *C. elegans*, although mutation of either the ubiquitin-activating enzyme (*uba-1*, E1), the ubiquitin-conjugating enzymes (*ubc* genes, E2s), the Skp1-related (*skr*) genes, or the Cullin (*cul*) genes results in a wide array of developmental phenotypes (Kipreos, 2005). Intriguingly, CUL-6, along with the redundant SKR-3, SKR-4, SKR-5 proteins, promote proteostasis during infection with *Nematocida parisii*, a microsporidia intracellular pathogen (Bakowski et al., 2014; Panek et al., 2020). In this context, SKR-5 is functionally the most important. We find a role for *cul-6* in vitellogenin regulation but only when *skr-3* is also deleted, suggesting that other Cullin proteins may act redundantly with CUL-6. Regardless, SKR-3 is functionally the most important of the SKR-3,4,5 family member for vitellogenesis. Thus, the intestinal SKR preference during the *N. parisii* infection differs from that during vitellogenesis, which in turn likely dictates the F-box adapter. Indeed, FBXA-75 and FBXA-158 function during *N. parisii* infection while FBXL-5 governs ubiquitin-mediated metabolic regulation (Panek et al., 2020).

In mammals, the mTOR signaling pathways are regulated by ubiquitination in several different contexts (Jiang et al., 2019). To our knowledge, FBXL-5 represents the first F-box protein to function in mTOR signaling outside of mammals, although it remains unclear whether FBXL-5 directly acts on a component of the mTORC2 complex. Mutation of *fbxl-5* largely mirrors the transcriptional response to loss of mTORC2; however, the *fbxl-5* mutation is unlikely to also impair mTORC1 activity, as loss of both complexes would be predicated to result in severe developmental defects. It is possible that FBXL-5 mediates switching of factors between the two mTOR complexes. This phenomena has been described in mammals, whereby the TRAF2 E3 ligase mediates K63-linked poly-ubiquitination of GβL/LST8, a shared component of both mTORC1 and mTORC2, which promotes mTOR-Raptor binding and dissociation of Sin1 from mTORC2, resulting in elevated mTORC1 signaling and reduced mTORC2 signaling (Wang et al., 2017). Alternatively, loss of FBXL-5 may directly impact the stability or activity of a core mTORC2 component; however, this hypothesis argues that ubiquitination would activate mTORC2 or that FBXL-5 inhibits a negative regulator of mTORC2 signaling. Consistent with the latter hypothesis, proteosomal degradation of Deptor, which itself inhibits both mTORC1 and mTORC2, promotes the activity of both complexes in vertebrates (Wang et al., 2017); however, a functional homologue of Deptor has yet to be identified in *C. elegans*. In the future, it will be crucial to identify the target of FBXL-5 to gain mechanistic insight into how ubiquitination of mTORC2 signaling components may govern intestinal homeostasis.

Loss of *fbxl-5* resulted in an increase in the transcription of most ribosomal protein genes. Interestingly, the expression of the cytosolic ribosomal proteins is reduced in the germline-less *glp-1(e2141)* mutant in a *daf-16*-dependent manner, suggesting that DAF-16/FOXO may be a transcriptional repressor of ribosomal protein gene expression (Hemphill et al., 2022). Moreover, reduced protein translation from impaired mTOR/DAF-16 signaling is likely to contribute to lifespan extension in multiple contexts (Hansen et al., 2007; Pan et al., 2007). It is therefore possible that the increase in ribosomal protein gene expression may contribute to the shorter lifespan that we observed in the *fbxl-5* mutant. Paradoxically, over-expression of *fbxl-5* in the intestine dramatically reduces lifespan and the *fbxl-5* loss-of-function mutation extends the lifespan of the *rict-1* mutant, suggesting that FBXL-5 may function in multiple pathways or in different tissues to regulate aging. Intriguingly, *fbxl-5* is up-regulated in the intestine of aging adults, which requires the homeodomain transcription factor UNC-62 (Van Nostrand et al., 2013). It is possible that a post-reproductive function of FBXL-5 may be to down-regulate vitellogenesis by simultaneously restricting vitellogenin transcription and translation by reducing the expression of the *vit* genes and ribosomal protein genes, respectively.

The FBXL-5 protein acts in a SCF complex to govern ubiquitin-mediated proteasome degradation of a component within the mTORC2 pathway. We find that levels of FBXL-5 are controlled by the hypodermal heterochronic pathway. Our study elucidates a new mechanism by which intestinal metabolic regulation is coupled to a developmental timing pathway. Conceivably, dynamic up-regulation of *fbxl-5* in response to adverse environmental conditions, which could include reduced food availability or pathogen infection, might tune vitellogenin expression levels to match the energetic resources that are available. Alternatively, FBXL-5 might purely act as a developmental regulator that responds to a variety of inter-tissue signals from different tissues; however, a systematic analysis of the signaling pathways that govern vitellogenesis will be needed to determine whether *fbxl-5* expression is under the control of additional developmental circuits. Importantly, identification of the target(s) of FBXL-5 could reveal a new upstream regulator of mTORC2 signaling or a novel ubiquitination-dependent mechanism of mTORC2 activation.

## DATA AVAILABILITY STATEMENT

The raw data for the mRNA-Seq expeiments has been deposited in the Gene Expression Omnibus (GSE248602). All additional raw data that support the conclusions described in this article will be made available by the authors, without undue reservation.

## AUTHOR CONTRIBUTIONS

R.H.D designed the experiments and P.C.B, K.G.K, M.A.N., and R.H.D performed the experiments and interpreted the results. R.H.D prepared the manuscript.

## ACKNOWLEDGEMENTS

The Caenorhabditis Genetics Center provided some of the strains used in this study and is supported by the NIH Office of Research Infrastructure Programs (P40 OD010440). We thank Yasmine Ackall and Monica Macharios for their assistance in carrying out some of the experiments described in this study.

