## Supplemental Material for "The F-box protein FBXL-5 governs vitellogenesis and lipid homeostasis in *C. elegans*"

**Contents** **Page**


1. Supplemental Tables S1-S3 2
2. Supplemental Figures S1-S4 6
3. References 10

| Strain | Genotype | Reference |
|---|---|---|
| N2 | Wild-type | (Brenner, 1974) |
| DLS258 | *lin-4(e912) II; fbxl-5(rhd43) V; mgIs70[*P*vit-3::GFP]* | This study |
| DLS260 | *lin-4(e912) II; fbxl-5(rhd56) V; mgIs70[Pvit-3::GFP]* | This study |
| DLS316 | *fbxl-5(rhd43) V* | This study |
| DLS327 | *lin-4(e912) II; fbxl-5(rhd43) V; mgIs70[*P*vit-3::GFP]; rhdEx73[*P*vha-6::mCherry::his-58::SL2::fbxl-5* + P*myo-2::mCherry]* | This study |
| DLS330 | *lin-4(e912) II; fbxl-5(rhd43) V; mgIs70[*P*vit-3::GFP]; rhdEx76[*P*col-10::mCherry::his-58::SL2::fbxl-5* + P*myo-2::mCherry]* | This study |
| DLS333 | *lin-29(n333) II; fbxl-5(rhd43) V* | This study |
| DLS344 | *rhdEx93[*P*fbxl-5::mCherry::unc-54 3'UTR* + P*myo-3::GFP]* | This study |
| DLS445 | *mgIs70[*P*vit-3::GFP]; rhdIs2[*P*vha-6::mCherry::his-58::SL2::fbxl-5 + cb-unc-119(+)]* | This study |
| DLS447 | *mgIs70[*P*vit-3::GFP]; rhdIs4[*P*vha-6::mCherry::his-58::SL2::fbxl-5 + cb-unc-119(+)]* | This study |
| DLS476 | *rhdIs2[*P*vha-6::mCherry::his-58::SL2::fbxl-5 + cb-unc-119(+)]* | This study |
| DLS477 | *rhdIs4[*P*vha-6::mCherry::his-58::SL2::fbxl-5 + cb-unc-119(+)]* | This study |
| DLS490 | *rict-1(mg360) II* | This study |
| DLS491 | *rict-1(mg360) II; fbxl-5(rhd43) V* | This study |
| DLS492 | *lin-29(n333) II; fbxl-5(rhd43) V* | This study |
| DLS537 | *rhdSi42[*P*vit-3::mCherry::unc-54 3'UTR + cb-unc-119(+)] II* | (Torzone et al., 2023) |
| DLS561 | *rhdSi42[*P*vit-3::mCherry::unc-54 3'UTR + cb-unc-119(+)] lin-4(e912) II* | This study |
| DLS708 | *cul-6(ok1614) IV; mgIs70[*P*vit-3::GFP]; rhdIs4[*P*vha-6::mCherry::his-58::SL2::fbxl-5 + cb-unc-119(+)]* | This study |
| DLS709 | *skr-3(ok365) V; mgIs70[*P*vit-3::GFP]; rhdIs4[*P*vha-6::mCherry::his-58::SL2::fbxl-5 + cb-unc-119(+)]* | This study |
| DLS726 | *cul-6(ok1614) IV; skr-3(ok365) V; mgIs70[*P*vit-3::GFP]; rhdIs4[*P*vha-6::mCherry::his-58::SL2::fbxl-5 + cb-unc-119(+)]* | This study |
| DLS806 | *cul-6(ok1614) IV; skr-3(ok365) skr-5(rhd269) skr-4(rhd283[W67*]) V; mgIs70[*P*vit-3::GFP]; rhdIs4[*P*vha-6::mCherry::his-58::SL2::fbxl-5 + cb-unc-119(+)]* | This study |
| DLS817 | *cul-6(ok1614) IV; skr-3(ok365) skr-5(rhd269) V; mgIs70[*P*vit-3::GFP]; rhdIs4[*P*vha-6::mCherry::his-58::SL2::fbxl-5 + cb-unc-119(+)]* | This study |
| DLS863 | *uba-1(it129) IV; mgIs70[Pvit-3::GFP]; rhdIs4[*P*vha-6::mCherry::his-58::SL2::fbxl-5 + cb-unc-119(+)]* | This study |
| DLS874 | *reSi5[*P*ges-1::TIR1::F2A::mTagBFP2::NLS::AID::tbb-2 3'UTR] I; rhdSi42[*P*vit-3::mCherry::unc-54 3'UTR + cb-unc-119(+)] II; fbxl-5(rhd298[3xFLAG::AID::fbxl-5]) V* | This study |
| DLS884 | *rhdSi42[*P*vit-3::mCherry::unc-54 3'UTR + cb-unc-119(+)] lin-4(e912) II; fbxl-5(rhd304) V* | This study |
| DLS885 | *rhdSi42[*P*vit-3::mCherry::unc-54 3'UTR + cb-unc-119(+)] lin-4(e912) II; fbxl-5(rhd305) V* | This study |

| | | |
|---|---|---|
| DLS886 | *rhdSi42[*P*vit-3::mCherry::unc-54 3'UTR + cb-unc-119(+)] lin-4(e912) II; fbxl-5(rhd306) V* | This study |
| DLS889 | *reSi5[Pges-1::TIR1::F2A::mTagBFP2::NLS::AID::tbb-2 3'UTR] I; rhdSi42[*P*vit-3::mCherry::unc-54 3'UTR + cb-unc-119(+)] lin-4(e912) II; fbxl-5(rhd298[3xFLAG::AID::fbxl-5]) V* | This study |
| DLS946 | *rict-1 & pqn-32(rhd314) II; mgIs70[Pvit-3::GFP]* | This study |
| DLS948 | *rict-1(mg360) II; fbxl-5(rhd43) V; mgIs70[Pvit-3::GFP]* | This study |
| DLS949 | *rict-1 & pqn-32(rhd314) II; fbxl-5(rhd43) V; mgIs70[Pvit-3::GFP]* | This study |
| GR2122 | *mgIs70[*P*vit-3::GFP]* | (Dowen et al., 2016) |
| GR2123 | *lin-4(e912) II; mgIs70[*P*vit-3::GFP]* | (Dowen et al., 2016) |
| GR2125 | *lin-29(n333) II; mgIs70[*P*vit-3::GFP]* | (Dowen et al., 2016) |
| GR2140 | *sgk-1(ok538) X; mgIs70[*P*vit-3::GFP]* | (Dowen et al., 2016) |
| GR2146 | *daf-2(e1370) III; mgIs70[*P*vit-3::GFP]/+* | (Dowen et al., 2016) |
| GR2147 | *rict-1(mg360) II; mgIs70[*P*vit-3::GFP]* | (Dowen et al., 2016) |
| MT333 | *lin-29(n333) II* | (Ambros and Horvitz, 1984) |

**Supplementary Table S1. *C. elegans* strains used in this study.** The strain names, genotypes, and associated references are shown.

| Target gene | Location in gene, crRNA guide number | crRNA sequence | Alleles | Genomic edit |
|---|---|---|---|---|
| *skr-5* | 5' end, rhd32<br>3' end, rhd33 | rhd32: UCUCAUAAAAAGGCCUGUAA<br>rhd33: GGGCAAAUUUGGUCUUGAAG | *rhd269* | 682 bp deletion |
| *skr-4* | Internal, rhd53 | rhd53: UCCUUGAGAAGAUUAUCACC | *rhd283* | W67* |
| *fbxl-5* | 5' end, rhd60 | rhd60: UACCUUUCAAAUUUCCAGAA | *rhd298* | 3xFLAG::AID |
| *fbxl-5* | 5' end, rhd60<br>3' end, rhd12 | rhd60: UACCUUUCAAAUUUCCAGAA<br>rhd12: UCCAAUUGGUCCACACUCUG | *rhd304* | 2,474 bp deletion |
| *fbxl-5* | 5' end, rhd60<br>3' end, rhd12 | rhd60: UACCUUUCAAAUUUCCAGAA<br>rhd12: UCCAAUUGGUCCACACUCUG | *rhd305* | 2,508 bp deletion |
| *fbxl-5* | 5' end, rhd60<br>3' end, rhd12 | rhd60: UACCUUUCAAAUUUCCAGAA<br>rhd12: UCCAAUUGGUCCACACUCUG | *rhd306* | deletion, unknown size |
| *rict-1* | 5' end, rhd9<br>3' end, rhd40 | rhd9: AAAUUUCAAUUUUCAGGCGA<br>rhd40: GAAAAUACUUAUAAAUGGAA | *rhd314* | 18,298 bp deletion |

**Supplementary Table S2. The crRNAs used in this study.** A list of the crRNA guides that were used in this study, including their target genes, their ribonucleotide sequences, and the alleles and genomic edits that were generated with each edit.

| **mRNA Target** | **Primer Sequence (5’ to 3’)** | **Reference** |
|---|---|---|
| *act-1* | F: GCTGGACGTGATCTTACTGATTACC<br>R: GTAGCAGAGCTTCTCCTTGATGTC | (Hoogewijs et al., 2008) |
| *vit-1* | F: GAGGTTCGCTTTGACGGATA<br>R: GGCTTCACATTCCTCGTTCT | (Ding and Grosshans, 2009) |
| *vit-2* | F: GACACCGAGCTCATCCGCCCA<br>R: TTCCTTCTCTCCATTGACCT | (DePina et al., 2011) |
| *vit-3/4/5* | F: CATGTGCACCATCGAAGAACTC<br>R: CCAATGTGGTTTCAATGACAAGTTG | (Dowen et al., 2016) |
| *vit-6* | F: TTCACCCAGAAGCCAGTTC<br>R: AGGATGGGAGGCAGTAGAC | (Dowen et al., 2016) |
| *fbxl-5* | F: GCCAAACACAATCCAGTTCAG<br>R: AGAAGTCCGAAATCCAAGTCC | This study |

**Supplementary Table S3. The RT-qPCR primers.** The sequences (5’ to 3’) of the qPCR primers used in this study, as well as any associated references.

A

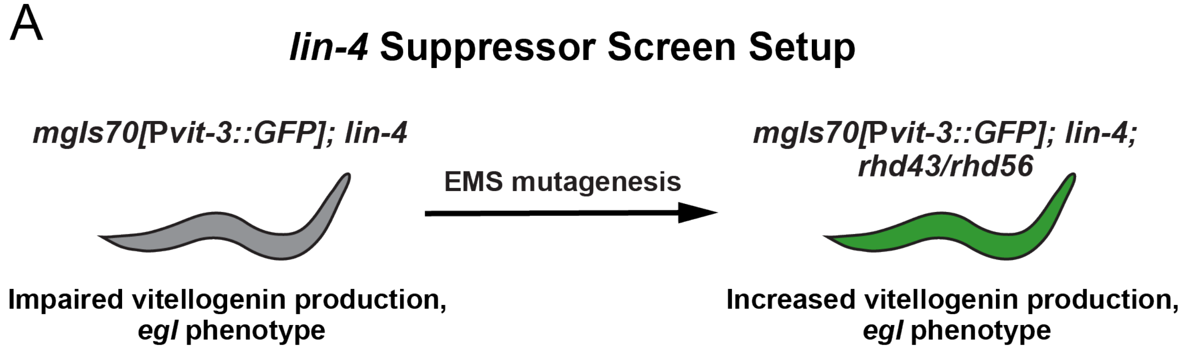


B

**T05B11.1/fbxl-5 Suppressor Alleles**

| Allele | Genetics | Change |
|---|---|---|
| *rhd43* | Recessive | Q113* |
| *rhd56* | Recessive | Splice Site Acceptor |

C

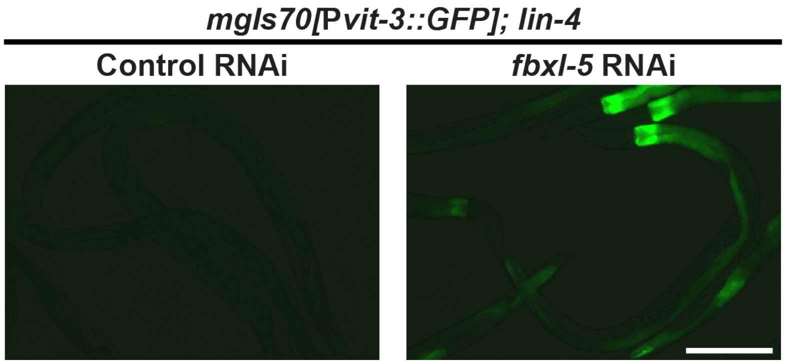


**Supplementary Figure S1. Mutations in the *fbxl-5* gene partially suppress the vitellogenesis defects displayed by the *lin-4* mutant. (A)** The design of the EMS mutagenesis screen employing the *mgIs70[*P*vit-3::GFP]* vitellogenesis reporter. The *lin-4(e912)* mutant fails to express the reporter, while the selected *lin-4* suppressor mutants express the reporter yet maintain the egg laying defects (*egl* phenotype) that are associated with the *lin-4* mutation. This approach selects for mutations that impact hypodermal-to-intestine developmental signaling and selects against mutations that act cell-autonomously in the vulva or hypodermis to suppress *lin-4* (*i.e.*, *lin-14* mutations). **(B)** The two *fbxl-5* mutant alleles isolated in the *lin-4* suppressor screen. **(C)** Representative fluorescence images of *mgIs70[*P*vit-3::GFP]* reporter expression in the *lin-4* mutant following control or *fbxl-5* RNAi (scale bar, 200 µm). The *mgIs70* transgene is an integrated high-copy transgene.

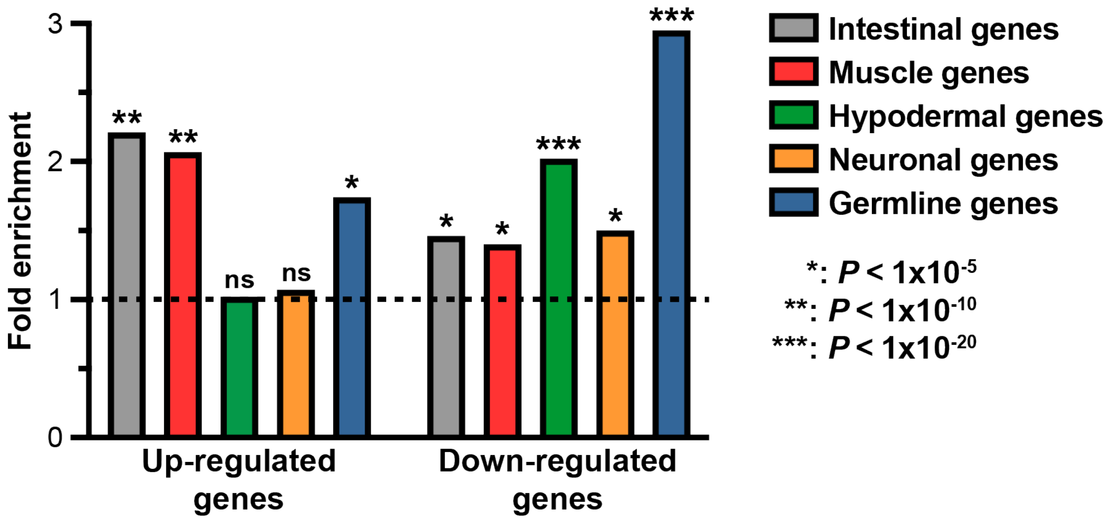


**Supplementary Figure S2. Loss of *fbxl-5* impacts gene expression in several different tissues.** Fold enrichment (observed/expected) for the differential expression of genes (mRNA-Seq of the *fbxl-5(rhd43)* mutant, 1% FDR) that are known to be expressed in the indicated tissues. The hypergeometric *P* values are reported.

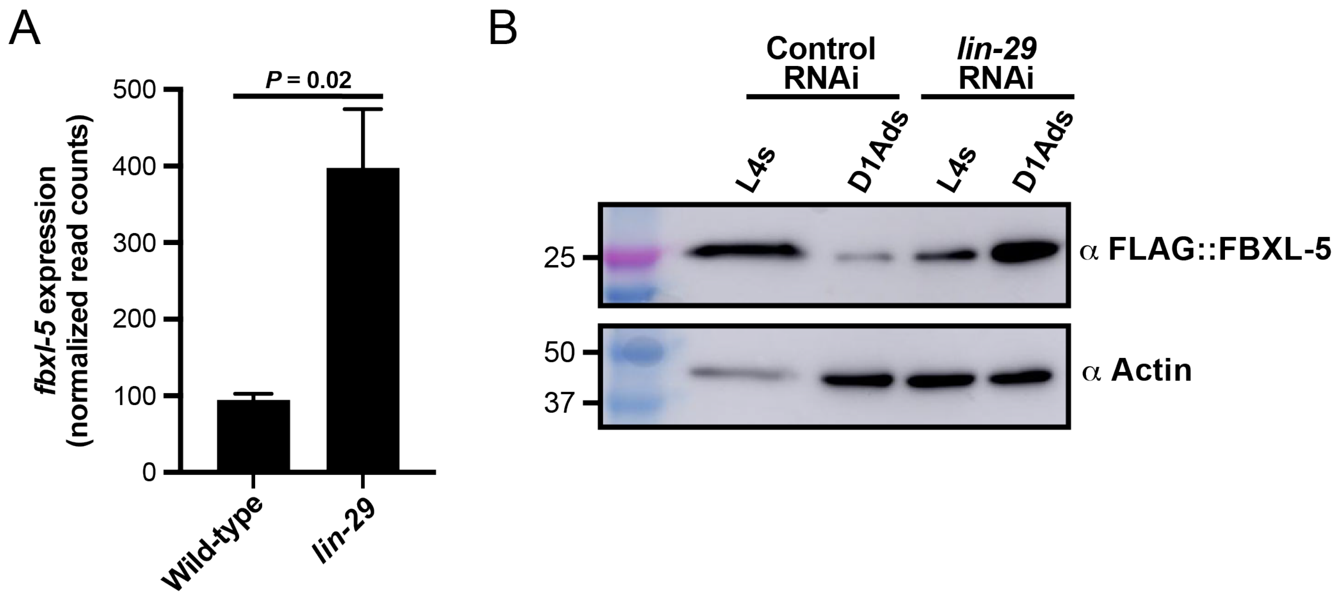


**Supplementary Figure S3. The *fbxl-5* gene is ectopically expressed in adult *lin-29(n333)* mutant animals. (A)** Normalized mRNA-Seq read counts for the *fbxl-5* gene in wild-type or *lin-29(n333)* mutant animals at the day 1 adult stage (mean +/- SEM; two-tailed T-test). **(B)** A western blot analysis of lysates from L4 or day 1 adult wild-type animals expressing an endogenously-tagged 3xFLAG::FBXL-5 protein after treatment with control or *lin-29* RNAi. An actin blot is included as a loading control.

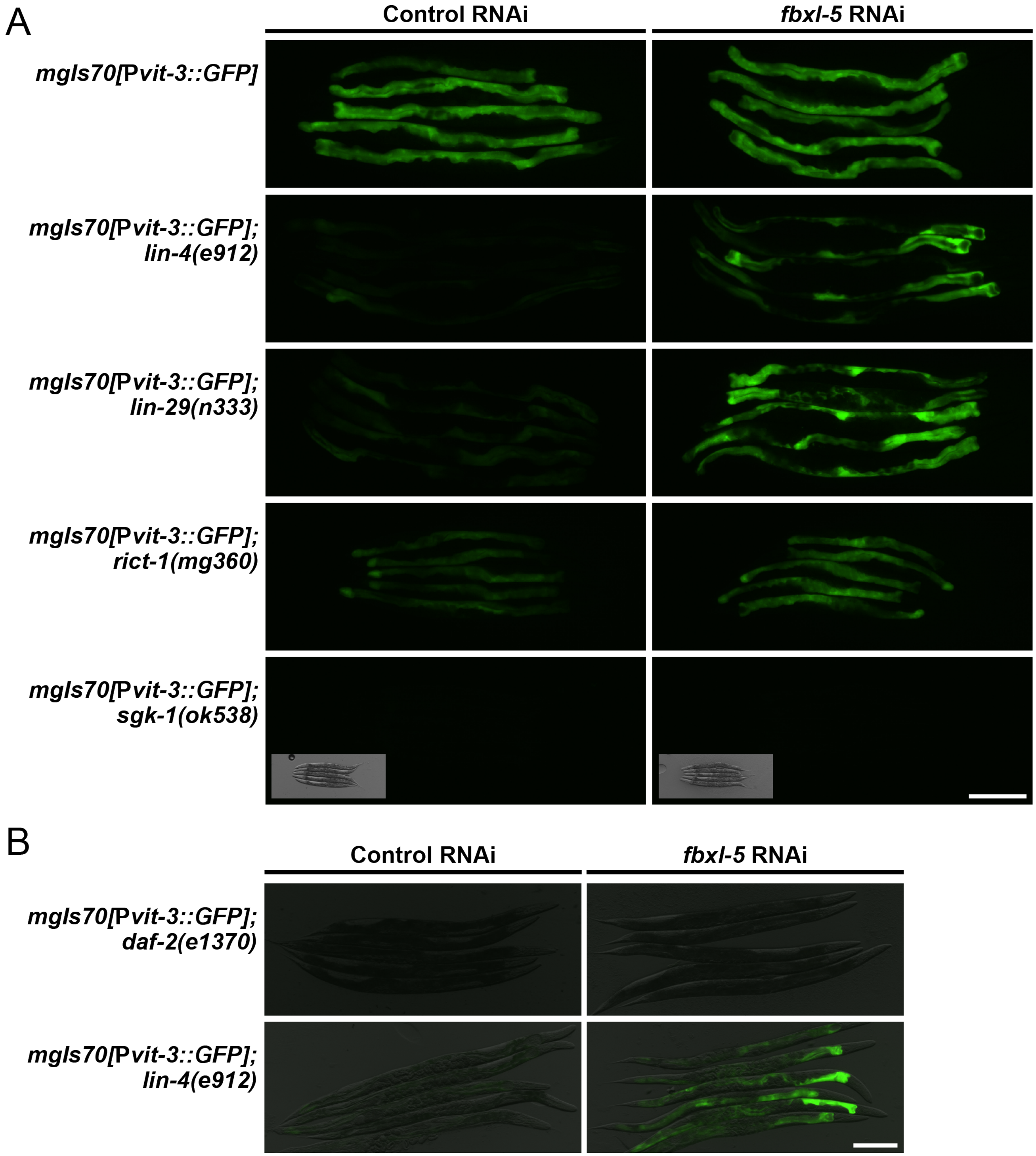


**Supplementary Figure S4. Knock-down of *fbxl-5* suppresses mTORC2, but not insulin, mutants.** **(A, B)** Representative P*vit-3::GFP* fluorescence images of the indicated mutants as day 1 adults following treatment with control or *fbxl-5* RNAi (scale bars, 200 μm). **(A)** The DIC images are included for the *sgk-1(ok538)* mutant since no GFP fluorescence is visible. **(B)** The GFP fluorescence images are overlaid on the DIC images.
